# Comparative transcriptomics reveals unique patterns of convergence in the evolution of eusociality

**DOI:** 10.1101/2021.07.11.451940

**Authors:** M. Velasque, Y. Tan, A.W. Liu, N.M. Luscombe, J.A. Denton

## Abstract

Eusociality represents a major evolutionary transition that arose independently in at least 12 insect lineages. Despite this prevalence, there remains considerable uncertainty surrounding the catalysing event and underlying genomic changes that enable such modifications. Commonly associated with this evolutionary transition is establishing and maintaining the reproductive division of labour (e.g. a reproductive queen and no-reproductive workers). This division is, at least in part, induced and maintained by highly species-specific pheromones. However, genomic analysis remains conflicted on the role of pheromones in this evolutionary transition. Specifically, if there was co-option of a common pheromone-sensitive genetic pathway present in all progenitor species or strong lineage-specific selection converging on similar transcriptomic signatures.

Using a solitary insect model, we sought to determine if various species-specific pheromones induced similar transcriptomic responses, thus activating similar pathways. We measured the transcriptomic and physiological response of a solitary insect, *Drosophila melanogaster*, to pheromones from bumblebees, honey bees, and termites. Each treatment induced the same strong physiological response - a decreased ovary size. However, employing several methods of transcriptomic analysis, we did not observe conservation in pheromone-mediated gene/pathway regulation.

Thus, despite a conserved phenotypic response, the underpinning transcriptome was vastly different. This suggests that pheromone-mediated eusociality is the result of convergent evolution. We propose that mechanisms maintaining eusociality (i.e. proto-pheromone) in early stages of eusocial evolution in each group, thus, acting as a primer for eusociality. This early state is then refined through strong selective pressure, resulting in a converging eusocial phenotype.

**Visual Abstract:** Figure 1.

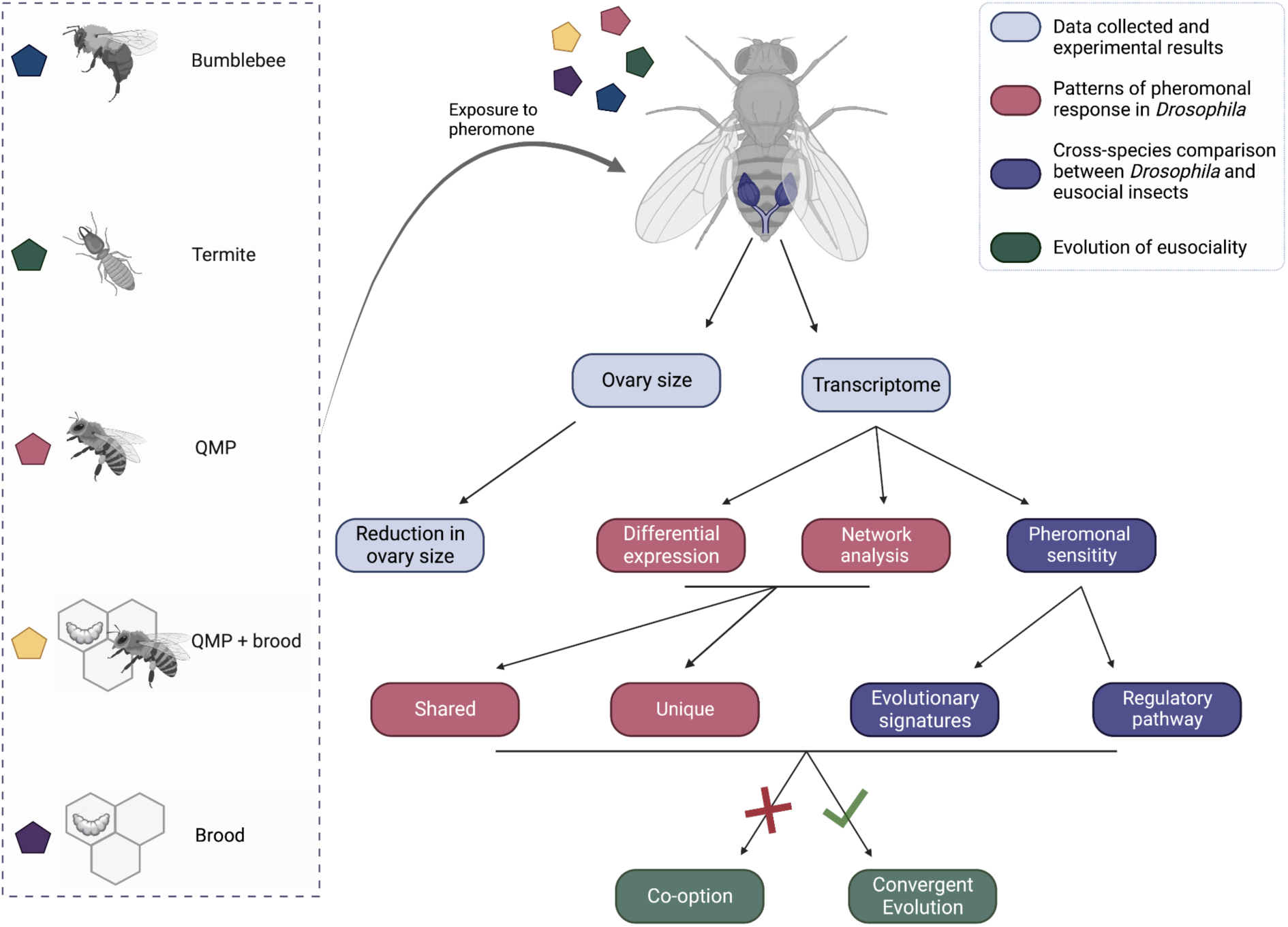

## Introduction

The transition from solitary to group living is a major evolutionary jump (1). However, it requires complex communication mechanisms to convey essential information to the group, such as the presence of predators and resource location (2, 3). Although there are multiple methods to transmit and receive information (e.g. visual, auditory, electric, olfactory), chemical cues are one of the most important and earliest to evolve (2, 4).

Insects use chemical components for recruitment, defence, predator recognition, and mate attraction, as well as to establish and shape social interactions (5–7). However, some eusocial organisms have evolved beyond context-dependent messaging. Acting as primer pheromones, they induce long-term behavioural and physiological effects by regulating colony structure and dynamics (8–11). This ranges from modulating aggression and attraction to monopolising colony reproduction through regulating female fecundity (12, 13). This monopolisation of reproduction by a few or one groups (queens/kings) creates a reproductive division of labour (8, 9). This is the cornerstone of eusociality (14–17).

Pheromone-mediated eusociality is not unique to honey bees as it has independently evolved at least 12 times in arthropods (18). The diverse evolutionary history of eusociality is also reflected in the production and structure of pheromones controlling the process (19–21). For instance, in several species, cuticular hydrocarbons (CHCs), a component ubiquitously produced in insects to prevent desiccation and infections, can act as queen pheromones (22–24). Such as in stingless and bumble bees, *Bombus terrestris,* where CHCs act as a queen pheromone (22, 25). In other species, such as the subterranean termite *Reticulitermes speratus,* queen pheromone is a mixture of CHCs, n-butyl-n-butyrate and 2-methyl-1butanol. However, the use of CHCs is not conserved in other eusocial animals. In *Apis mellifera,* queen pheromone comprises a complex blend of fatty acids secreted by the (26–28) mandibular gland (26–28). Moreover, pheromones produced by the brood (brood pheromone – Brood) were also found to regulate behaviour and worker reproduction in several species, such as *A. mellifera* (29–31).

In eusocial animals, the reproductive division of labour usually initiates during early larval development and is maintained throughout adulthood via exposure to queen pheromone (32, 33). Hence, in the absence of such pheromones, the division of labour breaks down as young workers activate their ovaries and engage in more selfish behaviour (34–38).

In eusocial animals, the division of labour and reproduction are suggested to be intertwined, arising from modifications of the reproductive life cycle of solitary insects (39, 40). Comparative genomics of multiple eusocial insect species corroborates this hypothesis and suggests that eusocial traits evolved from the co-option of pathways present in solitary ancestors (41–45). Moreover, there is strong evidence that specific queen pheromones evolved from conserved signals of solitary ancestors (22, 45, 46). It has been proposed that eusociality requires a highly conserved group of core genes, present in the solitary ancestor, that serves as a ground plan for eusociality (43). However, clear unambiguous evidence of this co-option hypothesis remains elusive (42, 47).

Further expansion of comparative genomics has added to this ambiguity. Genome analysis revealed branch-specific positive selection in five bee species from the *Apis* genus (48). Specifically, a greater proportion of species-specific genes were under positive selection than *Apis*-specific (45, 48). Furthermore, despite similar eusocial behaviour, three eusocial termite species demonstrated vastly different underpinning genetic mechanisms when contrasted against a wide diversity of non-eusocial and eusocial insect species (49). Thus, eusociality may be induced and maintained through diverse genetic means.

Pheromone-induced transcriptomic changes reflect extensive evolutionary optimisation of affected networks. Thus, cross-species transcriptomic comparisons are complicated due to the evolutionary distances between eusocial insect species, as inter-species gene orthologs may have vastly different functions (44, 50–52). The birth of novel genes and the expansion or contraction of gene families can make clear orthologue identification impossible and can be conflated with evolutionary forces (52–54). Therefore, differences or similarities in expression patterns between two highly divergent species can be highly misleading (55). One approach to overcome this problem is by using a single species to investigate transcriptomic changes induced by various pheromones (56–58). This permits the study of conserved and co-opted transcriptomic changes and identifies molecular signatures of these rare evolutionary events.

Here, we investigate pheromone-mediated gene expression in a solitary species. Using *Drosophila melanogaster*, we examine if primer pheromones, regulating eusociality, evolved from the co-option of similar pathways present in a solitary ancestor. Specifically, whether distinct pheromones have a similar transcriptomic and physiological profile across multiple instances on which they evolved, as previously suggested. We present a comprehensive transcriptome dataset set investigating gene expression associated with the reproductive division of labour. We show the variation in differential gene expression induced termite (*Reticulitermes speratus*), bumblebee (*Bombus terrestris*) and honey bee (*Apis mellifera*) queen pheromone and honey bee brood pheromone. We leverage this data set to perform an up to date and comprehensive comparison of differentially vs distinct (i.e. non-homologous) expressed genes associated with the maintenance of eusociality.

## Results

### Pheromone-induced Ovary Changes

We identified a significant change in ovary size across all pheromone-treated *D. melanogaster* (Repeated measures ANOVA F_7,180_, p < 0.0001; Figure 1; Tables S1 & S2). Although all pheromone treatments strongly inhibited ovary size, their effects varied in strength. Bumblebee and Termite pheromone had similar (p = 1) severe pheromone-mediated ovary reductions of 0.32 mm^2^ and 0.35 mm^2^ (mean; n=10), respectively (Figure 1; Table S1). Pheromone treatments from *A. mellifera* queen mandibular pheromone (QMP), Brood and the QMP and Brood mix pheromone induced similar reductions in ovary size (QMP and Brood vs QMP p =1; QMP and Brood vs Brood p =1; QMP vs Brood p = 0.064) of 0.51 mm^2^, 0.65 mm^2^ and 0.59 mm^2^ (mean; n=10) respectively (Figure 1; Table S1).

**Figure 1.**
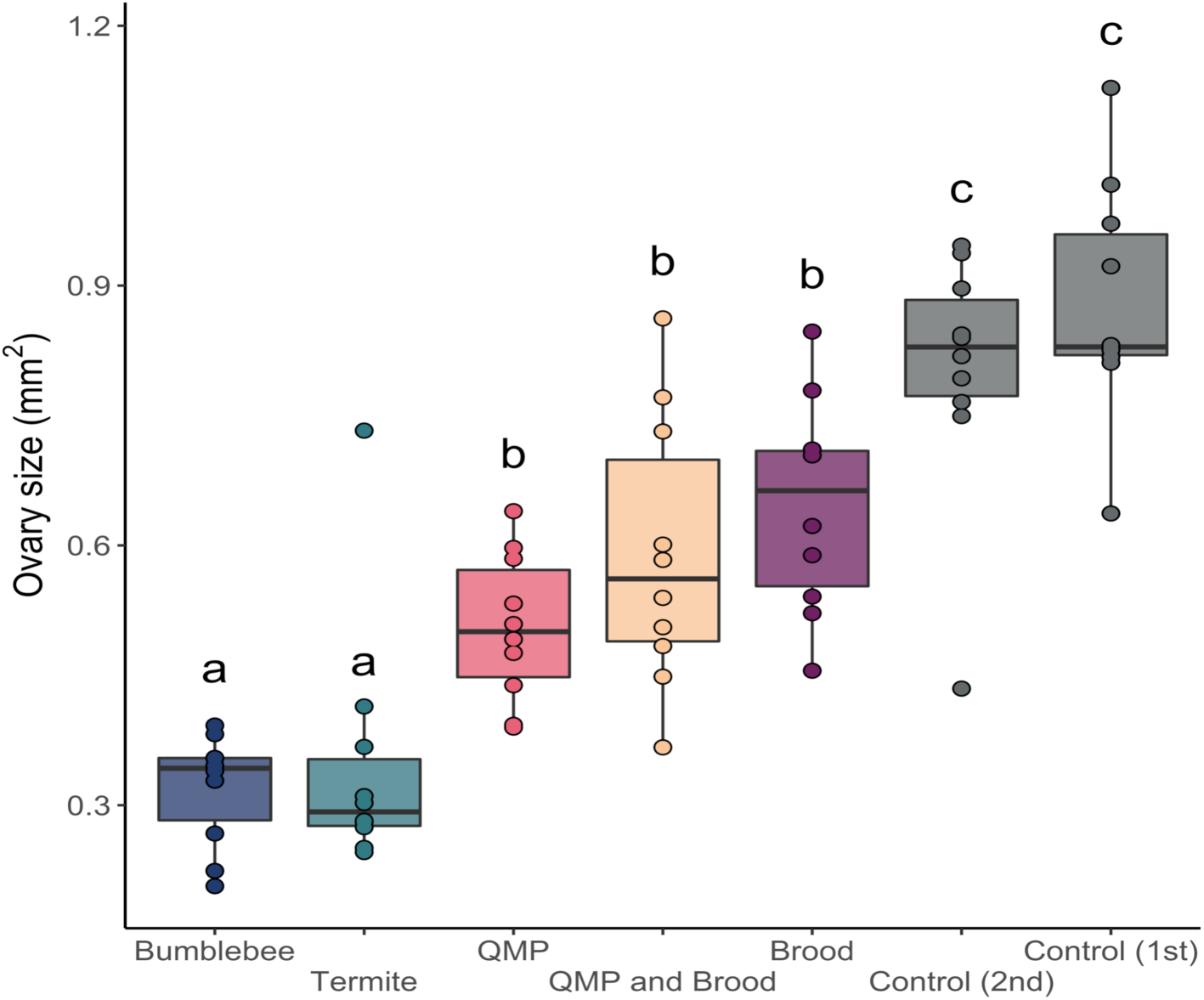
Pheromone treatment resulted in reduced *Drosophila melanogaster* ovary size. Treatments were queen pheromone produced by termites (*Reticulitermes speratus*), bumblebees (*Bombus terrestris*) and honey bees (*Apis mellifera*) (QMP), brood pheromone produced by honey bee larvae and a combination of both brood and QMP from honey bees. The pheromones were introduced to *D. melanogaster* from egg to adult via supplementation in food. Letters, either a, b or c, denote statistical significance with varying letters being significantly different (p-value < 0.05; Multiple Comparisons of Means: Tukey Contrasts; Table S2). Ten biological replicates were performed for each treatment or control with each biological replicate. Ovaries were measured from three flies per each replicate.

### Pheromone-induced Transcriptomic Changes

#### Differentially expressed genes and networks

Despite phenotypic similarities, we identified no transcriptomic overlap between pheromone treatment groups. Within each treatment, we identified similar numbers of differentially expressed genes with brood pheromone treatment being elevated: QMP (180 genes), Brood (640 genes), QMP and Brood (293 genes), Termites (207 genes) and Bumblebees (102 genes). There was no shared core of pheromone response genes and negligible inter-treatment overlap (Figure 2A; Figure S1). Due to their potential in eusociality regulation, differential expression of the Notch signalling pathway and the yellow gene family was specifically examined (59, 60). No differential expression of any genes encoding the 12 Notch signalling pathway core components or any of the 14 genes in the yellow family was detected in any treatment.

**Figure 2.**
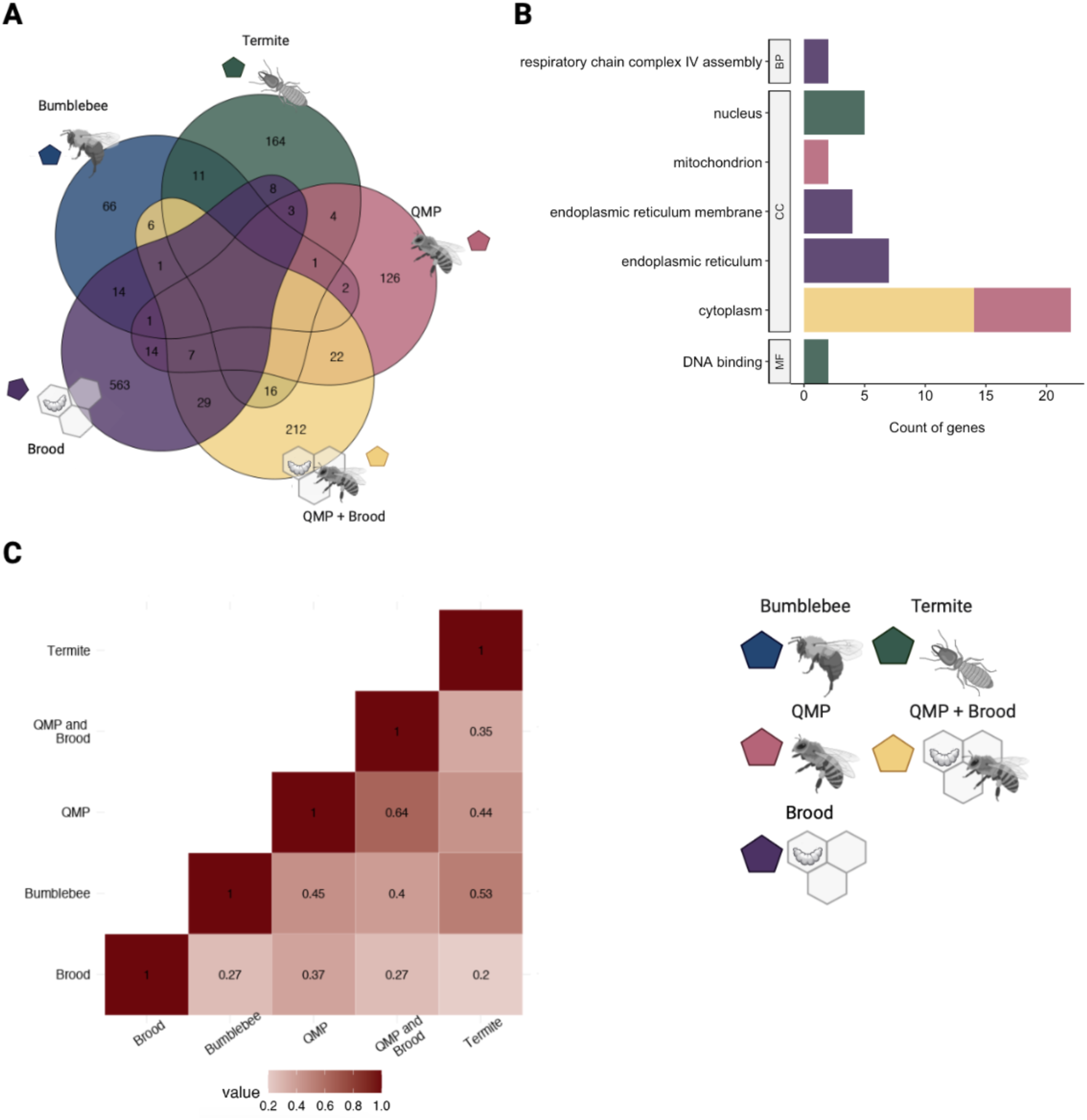
Differential gene expression of pheromone-sensitive pathways and genes in *Drosophila melanogaster* does not show a shared response. The DeSeq package was employed to determine differentially expressed genes in each of the pheromone treatments. A) Venn diagram showing the number of significantly differentially expressed genes per pheromone treatment. B) Results of signalling pathway analysis of differentially expressed genes across different pheromone treatments. C) Pearson’s correlation coefficient between the Log_2_-FC of significantly expressed transcripts in each treatment. Treatments were queen pheromone produced by termites (*Reticulitermes speratus*), bumblebees (*Bombus terrestris*) and honey bees (*Apis mellifera*), brood pheromone produced by honey bee larvae and a combination of both brood and queen pheromone from honey bees.

As we were unable to identify a core of differentially expressed genes, we investigated if pheromone-mediated changes in gene expression affected reproductive physiology through similar cellular processes. We compared gene enrichment of pheromone-induced changes in the transcriptome using GO (gene ontology) and KEGG (Kyoto Encyclopedia of Genes and Genomes) annotations. Although we identified some inter-treatment pathway enrichment, again, no core pathways were identified (Figure 2B).

In the absence of a conserved transcriptomic core, a broader general correlation between treatments was undertaken. There was a high degree correlation between the log_2_-fold-change (Log_2_FC) of all significantly expressed genes in each treatment (Figure 2C). To further investigate pheromone-mediated changes in genetic pathways and transcriptomic networks, we employed two different network analysis approaches.

#### Network Analysis

Employing network-based transcriptomic analysis, and weighted gene co-expression network analysis (WGCNA), no shared regulatory network common among all pheromone treatments was identified (Figure 3). We used WGCNA to screen and build a gene co-expression network, screen important modules, and identify candidate genes associated with each pheromone treatment (hub genes). We identified candidate genes by estimating the correlation between each gene and the pheromone treatment using Gene Significance analysis (61–63). WGCNA identified statistically significant differentially co-expressed genes between control groups and *D. melanogaster* treated with QMP (219 genes), Brood (219 genes), the mix of QMP and Brood (396 genes), Termites (307 genes) and Bumblebees (276 genes) (Figure 3). Once again, each treatment had limited overlap (Figure 3A; Figure S2). Employing GO enrichment and gene set enrichment analysis (GSEA) on the WGCNA differentially co-expressed genes also failed to identify high levels of overlap between functions (Figure 2B).

**Figure 3.**
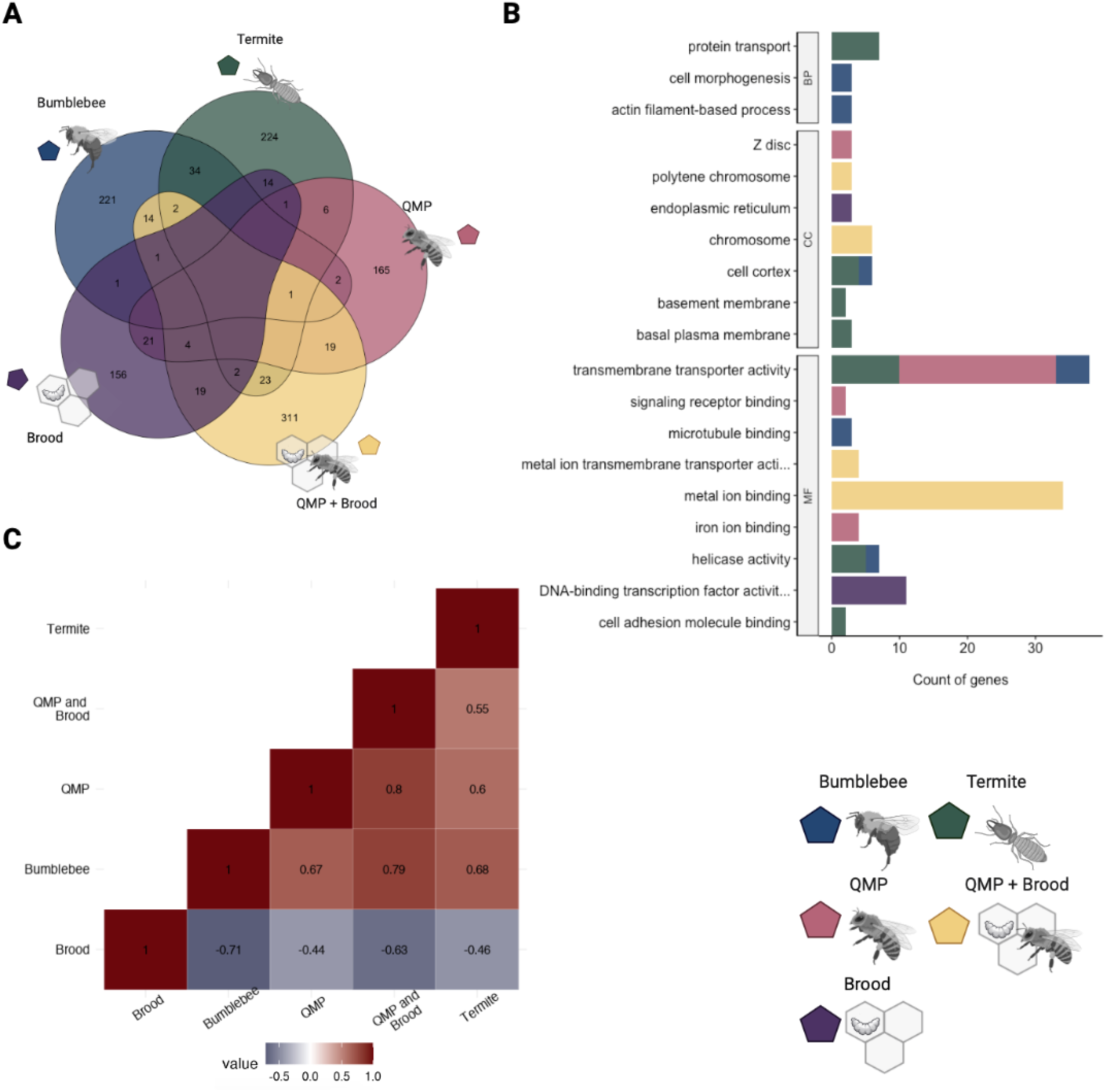
Network analysis showing pheromone-sensitive pathways and genes in *Drosophila melanogaster* does not show a shared response. Weighted gene co-expression analysis was used to screen and build a gene co-expression network, screen important modules, and identify candidate genes associated with each pheromone treatment. A) Venn diagram showing the number of significantly differentially expressed genes per pheromone treatment. B) Results of signalling pathway analysis of differentially expressed genes across different pheromone treatments. C) Pearson’s correlation coefficient between the Log_2_-FC of significantly expressed transcripts in each treatment. Treatments were queen pheromone produced by termites (*Reticulitermes speratus*), bumblebees (*Bombus terrestris*) and honey bees (*Apis mellifera*), brood pheromone produced by honey bee larvae and a combination of both brood and queen pheromone from honey bees.

Once again, there was a high degree correlation between the log_2_-fold-change (Log_2_FC) of all significantly expressed genes in each treatment (Figure 2C). However, Brood pheromone treatment resulted in a negative correlation for the WGCNA-identified transcripts. This was not observed in the mix of QMP and Brood treatment.

#### Cross-species comparison

We found that pheromone-mediated transcriptomic changes between eusocial insects and *D. melanogaster* were preserved. Although *D. melanogaster* has been previously used as a model organism in the study of eusociality (64), there are no direct comparisons between pheromone-mediated changes in gene expression for flies compared to eusocial insects. Holman et al. studied the transcriptomic response of several species, including *A. mellifera* and *B. terrestris,* treated with their own species’ pheromone (44). Using WGCNA module preservation, we determined the conservation of pheromone-mediated network signatures in our *D. melanogaster* experiments to their corresponding equivalents from Holman et al. Specifically, between our *D. melanogaster* exposed to QMP (*A. mellifera*) and Holman et al. *A. mellifera* exposed to QMP (*A. mellifera*), and our *D. melanogaster* exposed to bumblebee QMP (*B. terrestris*) and Holman et al. *B. terrestris* exposed to bumblebee QMP.

Comparing these datasets, we identified a high degree of WGCNA module preservation (Table 1; Figure 4; Figure S3). Overall, the *A. mellifera* comparison had a higher number of preserved modules and, thus, a greater number of shared genes in these modules. However, *A. mellifera* preservation also had a greater proportion of preserved modules with strong and moderate preservation (Figure 4A; Figure S3). The contrast between these two comparisons is likely down to the greater number of identified orthologues in *A. mellifera*. Regardless of the number, the preservation of these modules does demonstrate, in a broad sense, a similar transcriptomic response, and those serve as an external control for our analysis.

**Table 1.**
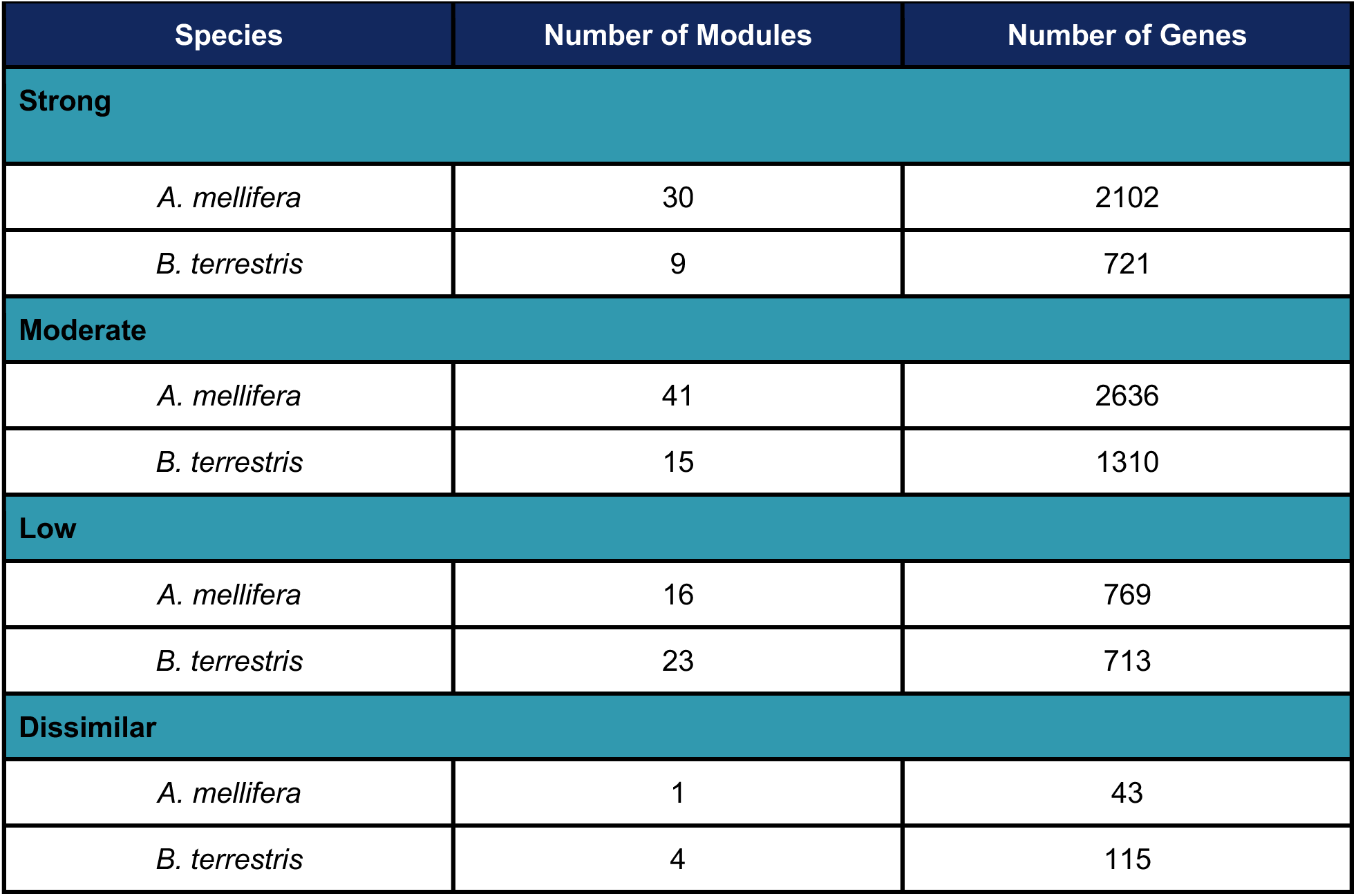
Cross-species model preservation of pheromone-response. The WGCNA module-preservation function facilitated comparisons between Drosophila melanogaster and Apis mellifera pheromone response to A. mellifera queen pheromone and D. melanogaster and Bombus terrestris pheromone-response to B. terrestris queen pheromone. Number of shared modules between the two. Total number of genes identified.

| Species | Number of Modules | Number of Genes |
| --- | --- | --- |
| **Strong** |  |  |
| *A. mellifera* | 30 | 2102 |
| *B. terrestris* | 9 | 721 |
| **Moderate** |  |  |
| *A. mellifera* | 41 | 2636 |
| *B. terrestris* | 15 | 1310 |
| **Low** |  |  |
| *A. mellifera* | 16 | 769 |
| *B. terrestris* | 23 | 713 |
| **Dissimilar** |  |  |
| *A. mellifera* | 1 | 43 |
| *B. terrestris* | 4 | 115 |

**Figure 4.**
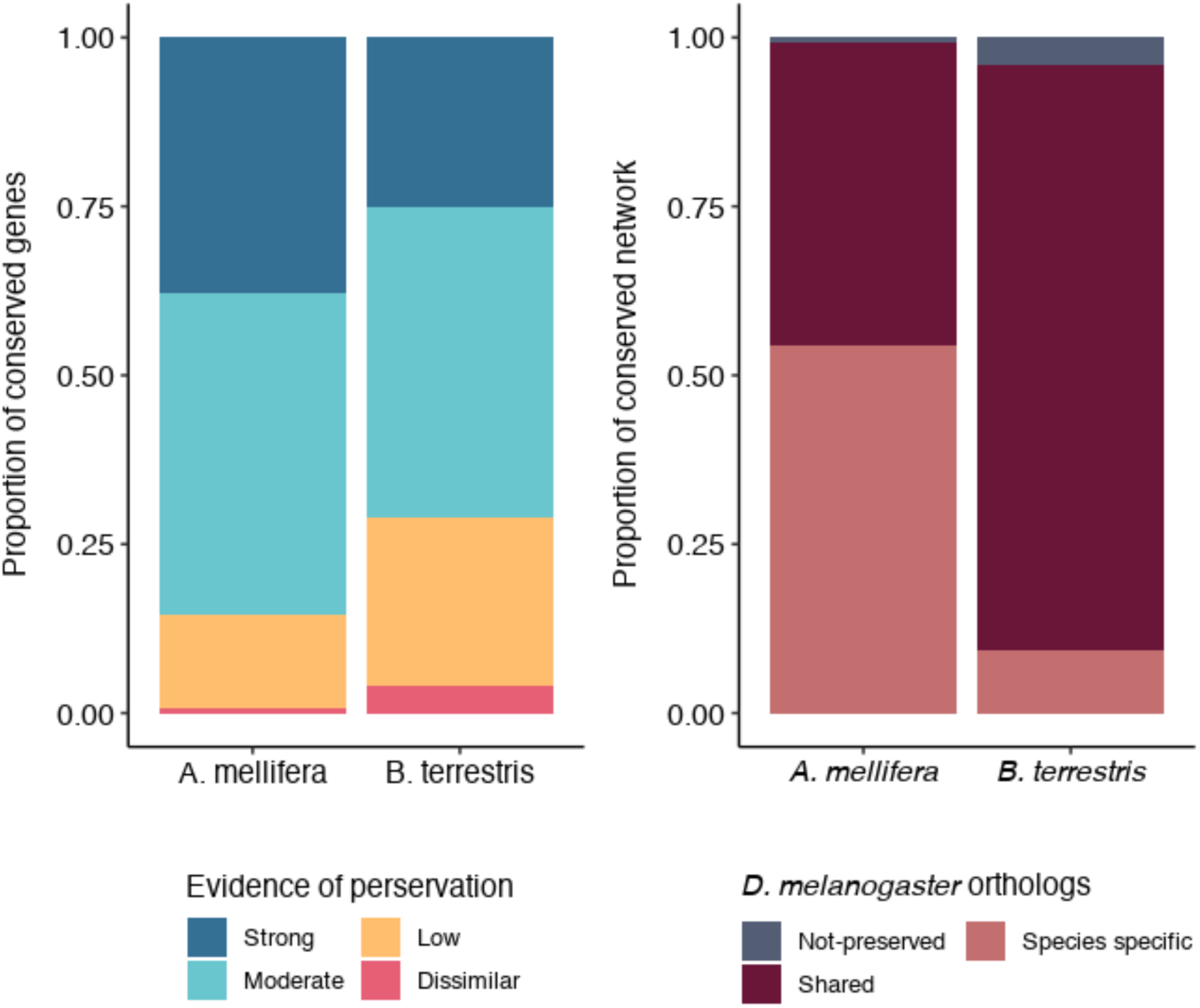
Pheromone-responsive module preservation between *Drosophila melanogaster* and *Apis mellifera* or *Bombus terrestris.* Comparing *D. melanogaster* pheromone-response data from this work and *A. mellifera* and *B. terrestris* pheromone-response data from Holman et al. Gene modules were determined using WGCNA module preservation. *A)* Proportion conserved pheromone sensitive genes in *D. melanogaster*, *A. mellifera* and *B. terrestris*. B) Summary of preservation evidence for pheromone response. Pheromonal sensitivity in *A. mellifera* and *B. terrestris* were compared with *D. melanogaster*. Shared genes indicate those highly conserved between the two comparison groups.

**Figure 4.**
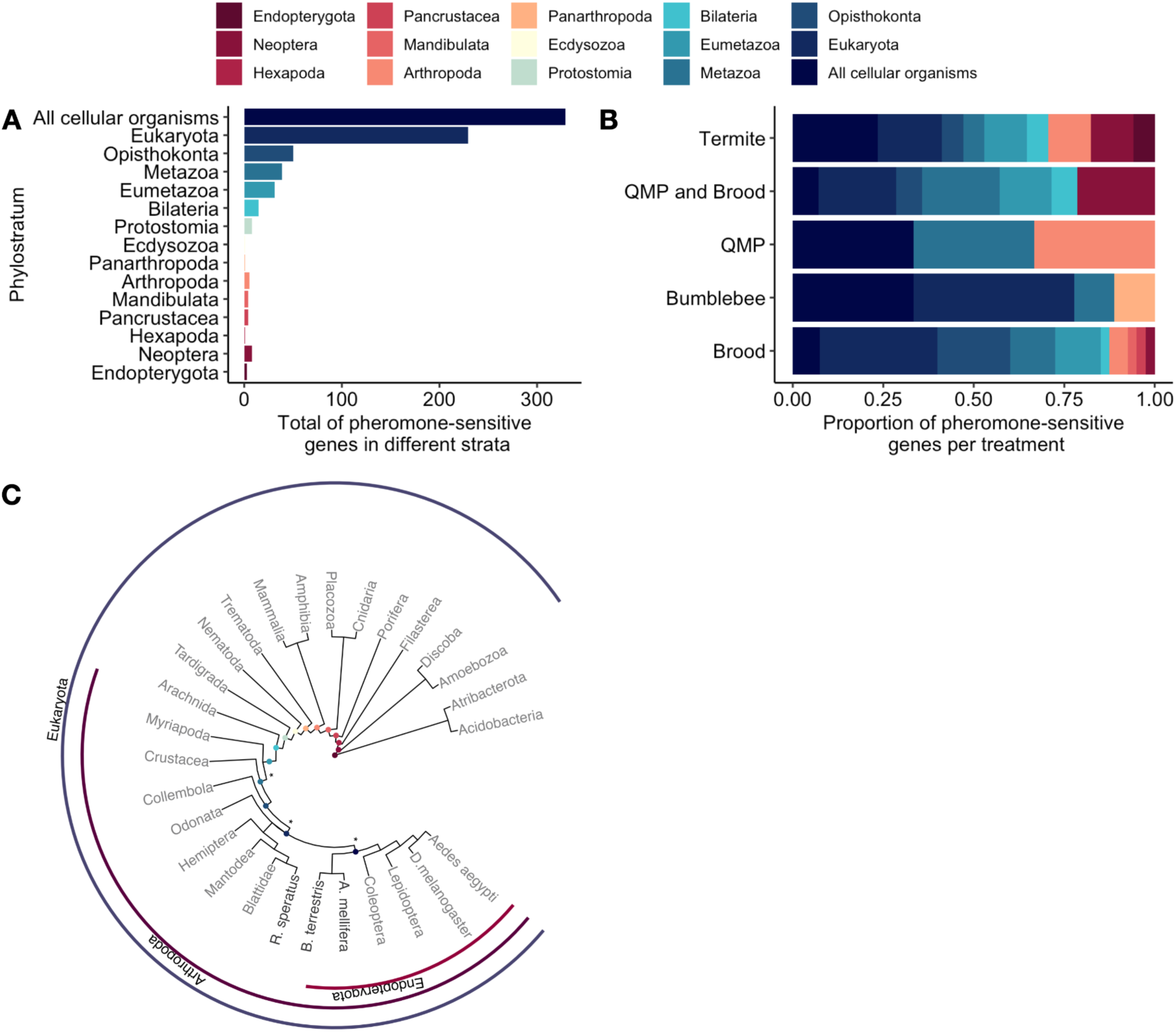
The evolutionary origins of pheromone responsive genes. The transcriptome age index (TAI) was used to infer the evolutionary origins of pheromone-responsive genes. A) Summary of all pheromone-sensitive genes per strata. B) Proportion of pheromone-sensitive genes in each treatment per strata. C) A phylogenetic tree with pylostratigraphic gene ages and phylogenetic relationships amongst 182 species. Dots indicate the evolutionary age of different pheromone-sensitive genes. Asterisks indicate phylogenetic levels (phylostrata) with significant changes in the expression of pheromone-sensitive genes.

#### Origins of Pheromone Response Genes

The responsive genes of each pheromone treatment do not have a conserved evolutionary origin, and thus eusociality is not co-oping recently evolved genes. The absence of overlapping differentially expressed pheromone-sensitive genes between treatments led to querying the evolutionary origins of these responsive pathways. We employed phylostratigraphy and calculated the transcriptome age index (TAI) to infer the evolutionary origins of pheromone-activated networks and genes. When separated by treatment, each group of differentially expressed pheromone-responsive genes lack a statistically significant conversed age (Figure 4A). If each treatment was further separated into up- and down-regulated differentially expressed transcripts, up-regulated transcripts from both termite and QMP treatments were statistically significant (Figure 4B).

A phylogenetic tree was created to determine the evolutionary relationship between target species homologues of the pheromone-responsive genes (Figure 4C). The tree was built using pylostratigraphic gene ages and phylogenetic relationships amongst a diverse collection of 182 species, with most species collapsed into systematic order or higher classification.

## Discussion

Although all pheromone treatments suppressed ovary development in *D. melanogaster,* we found no evidence of a shared evolutionary pathway or conserved mechanism underpinning this phenotype. In social insects, chemical communication has been independently co-opted in numerous species to facilitate the inhibition of worker reproduction by dominant females (4, 5). The prevailing hypothesis is that this induced sterility evolved from modifications in conserved molecular pathways regulating the physiology and behaviour of solitary species (44, 45, 59, 65, 66). Therefore, it was reasonable to expect similar pheromone-induced responses between different insect lineages and, thus, similar physiological and regulatory pathway responses. We removed the complexities of cross-species analysis and millions of years of convergent evolution by applying pheromones that inhibit worker reproduction from honeybees, bumblebees, and termites to a single species.

In contextualising our work, we are mindful of conflating factors introduced through cross-species analysis, especially when the species are separated by large evolutionary distances (18, 20, 48, 67). Furthermore, transcriptomic enrichment analyses, employed widely for cross-species studies, can potentially introduce biases (68, 69). However, we employed relaxed differential gene filtering (p<0.05, irrespective of expression fold-change) and diverse enrichment methods. Our view was that subsequent manual curation would allow us to identify candidate genes acting as universal eusocial catalysts. This proved unnecessary as we could not identify a core of differentially expressed genes and minimal inter-treatment overlap (Figure 2). Even GO enrichment and pathway analysis, highly permissive methods, only revealed minimal inter-group overlap.

Unable to identify a core of pheromone-responsive transcripts, we employed a candidate-based approach. Pheromone exposure has previously been shown to activate pathways related to lipid biosynthesis, oogenesis and olfaction (44, 45, 59). For instance, Notch appears to be an essential pathway regulating ovary development in the presence of queen pheromone (59) and it is generally assumed to be a co-opted pathway in solitary animals that facilitates eusociality (59). Although we found Notch to be differentially co-expressed in three of the five treatments, termite, QMP and QMP + brood, differing Notch signalling genes were affected in each treatment. An alternative candidate pathway is the yellow gene family. In Hymenoptera, this gene family is also responsible for regulating aspects of eusociality and includes a clade of genes referred to as the major royal jelly proteins (60, 70). In *D. melanogaster,* there are 14 members of this gene family (60, 71). The yellow gene family (*yellow-h* and *yellow-g2 in D. melanogaster*) was differentially co-expressed across all groups except for the brood treatment. Thus, these pathways seemingly provide a potential co-option target but do not appear to be a universal driver of pheromone response. Rather, given the widespread regulatory changes associated with eusociality (72–74), these gene families likely serve as candidate targets in refining eusocial traits.

Despite differences in the unpinning transcriptomic response, each treatment yielded smaller ovary size (Figure 1). As previous work demonstrated, pheromone exposure had no effect, even at much higher doses, on adult *D. melanogaster* (75). The ineffectiveness of pheromone-mediated reproductive inhibition on adults is also observed in other eusocial species (75–79). Insect oogenesis is a highly sensitive process disrupted by numerous environmental, developmental and gene-regulatory factors (80, 81). This includes the fact that cell death in the ovary is distinct from other tissues and mediated independently (80, 81). Furthermore, pheromone-mediated reproductive inhibition is not observed in primitively eusocial species like *Polistes satan* (paper wasp) or species with high reproductive plasticity like *Euglossa dilemma* (orchid bee) (41, 82). However, orchid bees form small 2–3 individual social groups wherein an abundance of specific cuticular hydrocarbons correlated with social dominance and ovary size (83). Therefore, oogenesis is a key candidate for pheromone-mediated disruption. Moreover, influencing oogenesis provides a potential mechanism for the monopolisation of colony reproduction and the evolution of eusociality.

Despite arising independently, several eusocial species share phenotypic and transcriptomic similarities (41–43). However, consistent with a growing body of comparative genomics (20, 84), we suggest this is not due to the co-option of identical or similar pathways in solitary ancestors. Rather, positive selection drives the convergence of characteristically eusocial phenotypes with diverse genomic origins. An example of this arises in the evolution of intra-colony communication. Insects rely on a host of olfactory signals and cues with the majority of these being detected by chemosensory receptors in the odorant receptor (OR) or ionotropic receptor (IR) gene families (85). These chemosensory pathways provide the core mechanism for intra-colony communication in several eusocial species (19, 20, 86, 87). Genomic analysis revealed that although termites and honey bees have expanded receptor gene families, termites had greater IRs and bees greater ORs (49, 86, 88–91). This means that although high-level analysis, such as GO, may detect these two gene family expansions as similar, closer examination demonstrated their independence.

*D. melanogaster* accurately depicts the effect of queen pheromone in eusocial species. When comparing our results with the transcriptome of honey bees and bumblebees exposed to queen pheromone (44), we found strong conservation of differentially expressed genes in pheromone treatments. Demonstrating the reliability of our results and the power of comparing the effect of distinct components (i.e. pheromones) in a single species.

Our work demonstrates the potential origins of eusociality evolution. The first steps of eusocial evolution likely began with a localised reproductive impediment that, over time, resulted in quasi-eusocial living. The absence of a core pheromone-induced transcriptomic response and yet conserved phenotypic response, suggests female fertility suppression can rapidly evolve using numerous compounds that induce regulatory changes. This is evident as eusociality has evolved at least eight times independently with limited overlap between the specific pheromones meditating this (67, 92).

The primer pheromonal signal in insects originated at least 100 Mya and evolved independently in multiple species. Their evolution and signal can be well traced across three groups: Blatoidea, Hymenoptera and Coleoptera, causing virtually a similar phenotype in all groups. The independent evolution of pheromones, the high conservation with their eusocial counterpart, combined with our results (i.e. pheromone treatment causing a similar ovary reaction in flies with no overlapping gene network and pathways) suggests pheromones are not a product of gene co-option, but the result of convergent phenotypic evolution. Producing a similar phenotype (inhibition of ovary development) through a different pathway in the species on which it evolved.

## Materials & Methods

We used the fruit fly *D. melanogaster* to investigate the hypothesis that eusociality evolved multiple times by co-opting existing regulatory pathways present in a solitary ancestor. We achieved this by studying the pheromone genetic signature of fruit flies exposed to pheromones from three different eusocial species: honey bee queen mandibular pheromone (QMP), bumblebee queen pheromone (Bumblebee), termite queen pheromone (Termite) and a pheromone produced by honey bee larvae, the brood pheromone (Brood). To simulate the conditions present in honey bee colonies, when they are exposed to both brood and queen pheromone, we also added a group with a mix of queen and brood pheromone (QMP + Brood).

### Fly husbandry

Flies were raised on standard Bloomington media at 24 °C under a 16h light / 8h dark cycle in 50 mL vials. The *D. melanogaster* strain Canton-S, a generous gift from the Van Vactor Unit, was used in all experiments. Each experimental vial contained ten males and females, of approximately 5-day-old. Each treatment group consisted of 10 vial replicates.

### Pheromone treatment

Bloomington fly food was made and cooled to >50°C before mixing in each phenomenon treatment (Table 2) at a 1 in 150 ml concentration. Ten millilitres of supplemented food were added to each vial and left to dry for 24 hours. Vials were kept at 25°C and under a powerful extractor to prevent cross-contamination due to pheromone volatility. Adult flies were left on the vial for four days to lay eggs and then removed and discarded. Virgin females were collected and allowed to mature for five days in vials containing the aforementioned food with the corresponding treatment pheromone.

**Table 2.** Pheromone Treatments, suppliers, and dilution.

| Treatment | Pheromone Formula | Supplier (ID) | Dilution |
| --- | --- | --- | --- |
| Control | - | - | 50 µl isopropanol |
| QMP (*Apis mellifera*) | 9-ODA, two isomers of 9-HDA, HOB and HVA | TempQueen (DC-705) | 20 µl of synthetic QMP mix diluted in 30 µl isopropanol |
| Brood pheromone (*Apis mellifera*) | Methyl palmitate, Ethyl palmitate, Methyl stearate, Ethyl stearate, Methyl oleate, Ethyl oleate, Methyl linoleate, Ethyl linoleate, Methyl linolenate, Ethyl linolenate | Contech (SuperBoost) | 10 µl SuperBoost diluted in 40 µl isopropanol |
| QMP and Brood pheromone mix (*Apis mellifera*) | 9-ODA, two isomers of 9-HDA, HOB and HVA; Methyl palmitate, Ethyl palmitate, Methyl stearate, Ethyl stearate, Methyl oleate, Ethyl oleate, Methyl linoleate, Ethyl linoleate, Methyl linolenate, Ethyl linolenate | TempQueen (DC-705) and Contech (SuperBoost) | - 20µL of synthetic QMP, 10µL Supperboost and 20µL isopropanol |
| Bumblebee queen pheromone (*Bombus terrestris*) | CHC pentacosane (C25) | Sigma-Aldrich (286931) | 1 µg C25 diluted in 50 µl isopropanol |
| Termite queen pheromone (*Reticulitermes speratus*) | n-butyl-n-butyrate and 2-methyl-1-butanol | n-butyl-n-butyrate Sigma-Aldrich (281964); 2-methyl-1-butanol Wako Chemicals (133-08385) | 15 µl n-butyl-n-butyrate and 7.5 µl 2-methyl-1-butanol diluted in 27.5 µl isopropanol |

### Ovary measurements

Ten biological replicates were used for each treatment or control. For each replicate, ovary collection for measurement was performed in triplicate and on the same day to prevent temporal changes in physiological measurements of ovaries. Ovary image identities were encoded such that they could be measured blind to reduce bias. Measurements were performed using ImageJ2 (93). We used normal QQplot and Shapiro-Wilk test to identify deviations from normality. Although termites had two outliers, all the points fall approximately along the reference line, and thus we assumed normality. We compared differences in ovary size between different treatment groups using One-way repeated-measures ANOVA.

### RNA Sequencing

For RNA sequencing, we randomly selected four vials and pulled five ovaries per vial. RNA extraction was performed with an in-house modified protocol (Supporting Text). Libraries were prepared using NEBNext® Ultra™ II Directional RNA Library Prep Kit (E7760L) for Illumina as instructed by the manufacturer.

### Data Analysis

The complete analysis pipeline and associated results are publicly available at https://github.com/marivelasque/eusociality_evolution. This pipeline is a modified version of our previously developed pipeline (94). RNA data were trimmed with Trimmomatic 0.38 (95) and quality was assessed using FASTQC (96). Transcript abundance was calculated with RSEM/bowtie2 (97, 98). Mapping and abundance calculations were performed against the *D. melanogaster* genome assembly BDGP6 (release 89). Differential expression analysis was performed using edgeR (99). We adjusted p-values to control the false discovery rate using the Benjamini-Hochberg method. The threshold of significance was defined by p<0.05. Visualisation relied on packages venn (100), ggplot2 (101), igraph (102) and ggraph (103). Analysis, figures and HTML files were generated using R in the RStudio environment.

### Gene co-expression network analysis

To investigate changes in gene networks mediated by eusocial pheromones we used Weighted Gene Co-Expression Network Analysis WGCNA (62). We used the WGCNA R package to construct the co-expression network using gene counts obtained from RSEM. Prior to the analysis, we removed gene outliers using the “goodSamplesGenes” (62) function and accessed overall sample clustering in the expression matrix using “hclust” (104). We used the lowest threshold power that resulted in approximate scale-free topology to construct the gene co-expression network using the blockwiseModules function (62). Then, we estimated the gene expression profile, module eigengene, and the module-treatment relationship, calculating the correlation between the module eigengene and treatment group. We implemented the analysis in two steps (i.e. lowest threshold power and differential co-expression) using the functions SoftPower and wgcna.est from (94). The threshold of significance was defined as p<0.05.

### Enrichment and pathway analysis

We obtained Gene Ontology (GO) and KEGG (Kyoto Encyclopedia of Genes and Genomes) annotations using the package biomaRt (105) clusterProfiler (104) respectively. We used Fisher’s exact test to identify significantly enriched genes using the package topGO (106) with the threshold of significance defined by P<0.05. We identified enriched pathways using Pathway-Express, a package part of the Onto-Tools (107). The significant pathways for predicting target genes were identified according to the Kyoto Encyclopedia of Genes and Genomes (KEGG) database obtained using the package clusterProfiler (104, 108). A minimal size of 40 genes of each geneSet was used, and the significance threshold was defined by p<0.05. Although there was an overlap on “Ribosome biogenesis in eukaryotes”, the enrichment showed a different directionality, overrepresented in Bumblebee, Termites, QMP, a mix of QMP and Brood over-represented in Brood pheromone.

The pathway analysis was implemented using the Pathway-Express package part of the Onto-Tools (107). Signalling pathways were identified using only differentially expressed genes. The relationship between each gene was inferred based on their KEGG pathway and magnitude of expression change (logFC,) and the visualisation of significantly enriched pathways was performed using Igraph (102).

### Cross-species comparison

To ensure pheromonal-induced changes in the fruit fly transcriptome are comparable to eusocial insects, we compared the network conservation of eusocial insects and fruit flies exposed to pheromone. Holman et al. (44) compared the whole transcriptome of several species, including *Apis mellifera* and *Bombus terrestris,* treated with either solvent-only control or their own species’ queen pheromone. To determine whether the QMP network signature was well conserved between *D. melanogaster*, *A. mellifera* and *B. terrestris,* we used a WGCNA integrated function (modulePreservation) (62) to calculate module preservation statistics. Orthologs between these two groups were obtained using BLAST-based on the *D. melanogaster* genome. Holman et al. data were obtained from BioProject DRP004516 (44). A total of 5507 unique *D. melanogaster* orthologs were identified in *A. mellifera* and 2744 in *B. terrestris*. To ensure that the conserved genes between the two data sets correspond to pheromone-sensitive genes, we compared the differential expression between each QMP treatment and control using exactTest function in edgeR (99). The threshold of significance was defined by P<0.05. No publically available pheromone termite treatment data were identified.

### Evolutionary Origins

We performed a protein-based homology search across the tree of life to determine the evolutionary age of pheromone-sensitive genes in *D.melanogaster* using Phylostratr (109). We compared *D. melanogaster* gene sequences against 182 spp. (NCBI proteins database) by the blastp algorithm at the e-value cutoff of 1e-03 and selected the best match for pheromone-sensitive ortholog in each species. Following, we compared the phylogenetic age of the pheromone-sensitive genes within each treatment using the Transcriptome age index (TAI)(110, 111). To estimate TAI, we retrieved the phylostrata (15 out of 27 phylostrata) of all genes that were differentially expressed across all pheromone treatments (all pheromone-sensitive genes) and their respective normalised expression. We used the pheromone-sensitive phylostrata to reconstruct phylogeny using ape and ggtree (112–115).

## Declarations

## Acknowledgements

The Okinawa Institute of Science & Technology Sequencing Centre kindly generated the sequencing data. Resources and expertise from the Okinawa Institute of Science & Technology High-Performance Computing Section were employed for data analysis. This work was funded by the Japanese Society for the Promotion of Science KAKENHI grant scheme (grant numbers 19K16205, 19K06795 awarded to MV and JAD, respectively) and the Okinawan Institute of Science & Technology. All authors were supported by the Okinawa Institute of Science & Technology. MV and JAD were also supported by Monash University and the World Mosquito Program Ltd, respectively.

## Availability of Data & Materials

All code and compiled markdown file showing will be available on GitHub prior to publication https://github.com/marivelasque/eusociality_evolution. A permanent copy of the final analysis markdown file is available at XXX <DOI provided prior to final publication>. All raw sequencing data are available under the BioProject XXX <PROVIDED prior to final publication>.

## Author Contributions

MV and JAD conceived the project, designed and performed the experiments, analysed the results and wrote the paper. YT, MV and AL prepared the sequencing libraries. All authors edited the manuscript and interpreted the results.

## Supporting Text & Materials

### Materials & Methods

#### RNA Extraction

A modified TRIzol RNA extraction protocol used for this work.

##### Materials

- 1 ml TRIzol
- 200 µl RNase-free chloroform
- 1 µl Glycogen
- 1 ml Isopropanol 100% w/w – chilled to -80°C
- 3 ml Ethanol 75% w/w – prepared fresh
- 500 µl Back Extraction Buffer (BEB)
- 20 µl Nuclease-free water

##### Sample homogenization and RNA extraction

1. Flash freeze samples (samples must be kept at -80oC) and transfer to dry ice (for processing) or -80oC freezer.
2. Add 500μL of TRIzol to each Eppendorf tube (keep samples in dry ice)
3. Homogenise sample with a pestle
4. Add 500μL of TRIzol
5. Mix by pipetting samples
6. Let it sit at room temperature for 5 min
7. Add 200μL of RNase-free chloroform (make sure it is a new vial or that it is only used on RNA work)
8. Shake vigorously by hand for 15s DO NOT VORTEX SAMPLES (vortex degrades the DNA)
9. Let it sit at room temperature for 3 min
10. Centrifuge at 4oC < 12,000g for 15 min
11. Transfer the upper aqueous phase to a new tube (leave 8-10μL in the bottom of the Eppendorf tube to prevent TRIzol contamination from getting into the next phase
12. Transfer the upper aqueous phase to a new tube (leave 8-10μL in the bottom of the tube to prevent TRIzol contamination from getting into the next phase. DO NOT DISCARD LOWER PHASE
13. Keep both upper and lower phases in separate tubes and proceed to 1-RNA and siRNA purification (upper aqueous phase) or 2-DNA purification (lower phase)

##### RNA and siRNA purification (upper aqueous phase)

1. Add 1μL of glycogen (can be skipped)
2. Add 500μL of RNase-free 100% isopropanol/ethanol (chilled to -80oC)
3. Invert by hand 10-20 times to mix (or very gently with the pipet)
4. Move samples to the fridge and let them rest overnight at -80 fridge
5. Centrifuge at 4oC < 12,000g for 10 min to precipitate the RNA (move samples very carefully not to agitate the mix, if disturbed, leave them overnight again).
6. Remove the supernatant and discard the rest.
7. Add 1μL of 75% Ethanol to pellet
8. Centrifuge at <7,500g for 5 min at 4oC
9. Remove supernatant and discard
10. Add 1μL of 75% Ethanol to pellet
11. Centrifuge at <7,500g for 5 min at 4oC
12. Remove supernatant and discard
13. Pulse spin samples at room temperature
14. Carefully remove the remaining supernatant with a pipette without disturbing the RNA pellet (use gel loading tips)
15. Air-dry the pellet at room temperature for 2-5 min to evaporate Ethanol
16. If necessary, heat tubes at 42°C to evaporate remaining Ethanol
17. Add 20μL of nuclease-free water to the pellet (over-dried RNA might take longer to dissolve)
18. Gentle mix the sample to solubilise the RNA
19. Place the solubilised RNA on ice immediately
20. Aliquot 8μL of the sample for quantification and cDNA synthesis (keep the remaining 12μL in - 80oC)
21. Quantify RNA concentration and purity.

**Figure S1.**
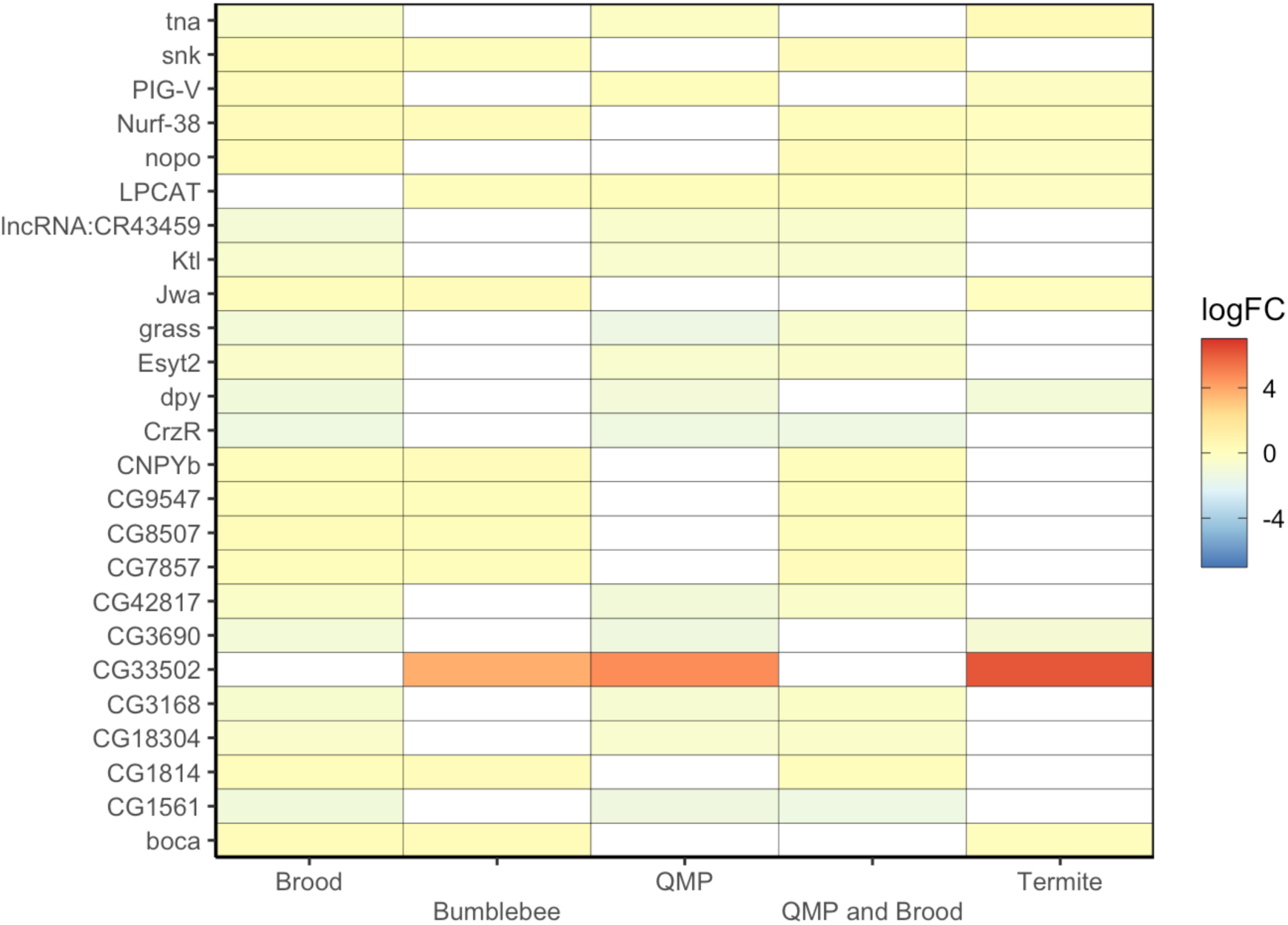
Log_2_–fold change heatmap of the top 25 differentially expressed genes. Differential gene expression was determined using deseq. Both up- and down-regulated genes were considered. The colour scale reflects the fold change in gene expression in pheromone compared to the controls, ranging from down-regulated (blue) to up-regulated (red). Threshold of significance was defined as p<0.05.

**Figure S2.**
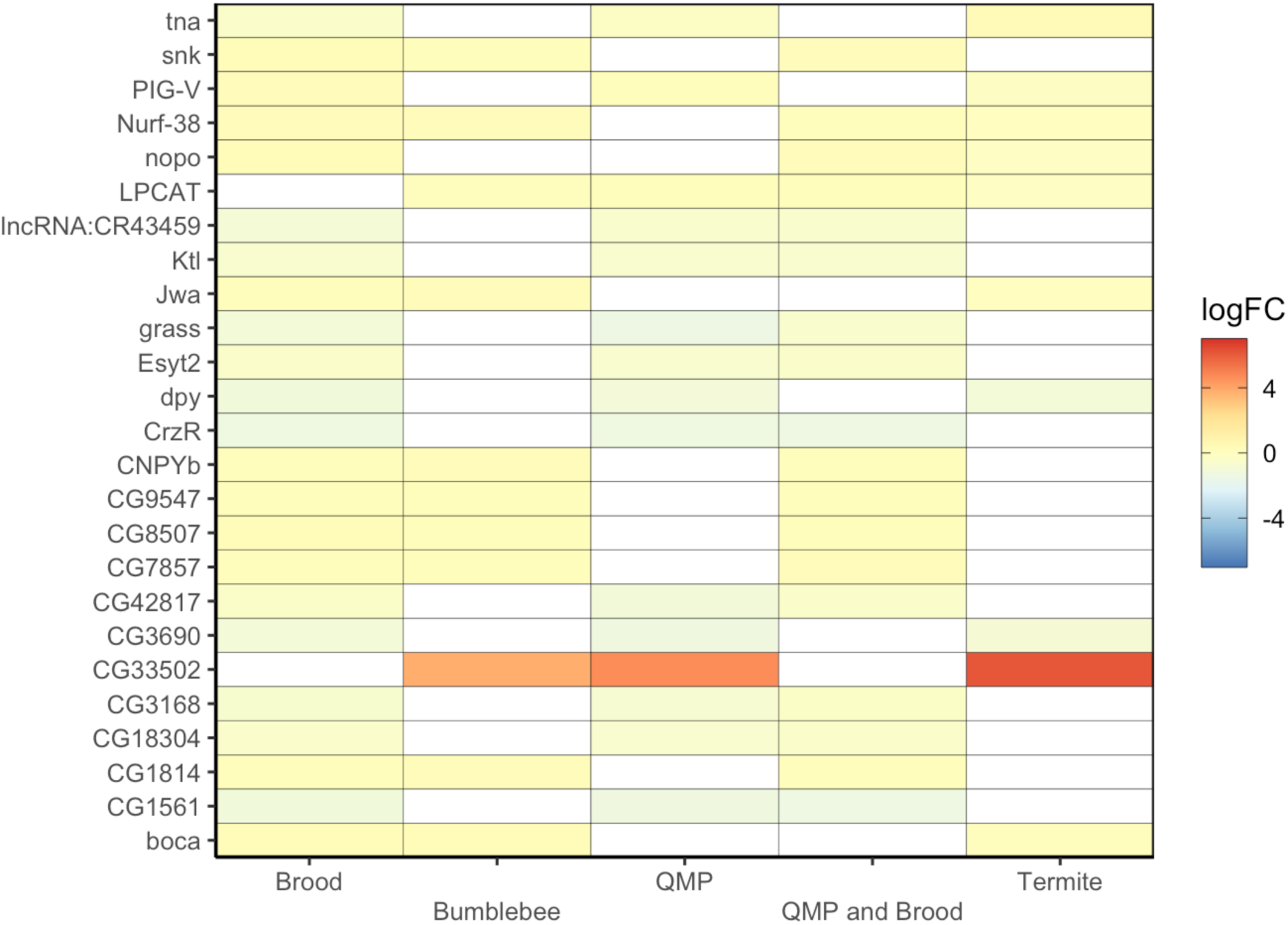
Log_2_–fold change heatmap of the top 25 differentially expressed genes. Differential gene expression was determined using WGCNA. Both up- and down-regulated genes were considered. The colour scale reflects the fold change in gene expression in pheromone compared to the controls, ranging from down-regulated (blue) to up-regulated (red). Threshold of significance was defined as p<0.05.

**Figure S3.**
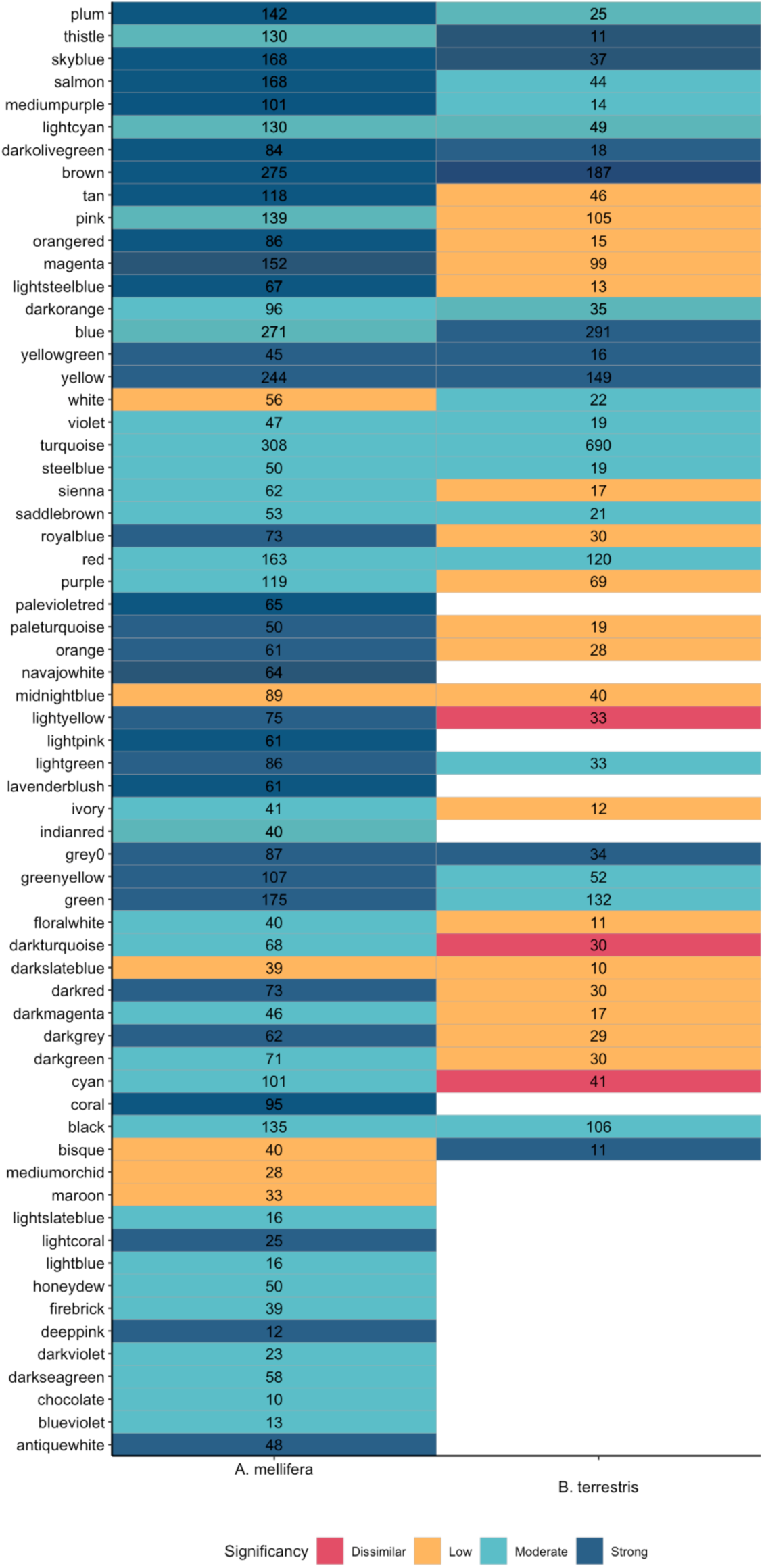
All pheromone-responsive modules preserved in cross-species comparison between *Drosophila melanogaster* and *Apis mellifera* or *Bombus terrestris.* The WGCNA module-preservation function facilitated comparisons between *Drosophila melanogaster* and *Apis mellifera* pheromone-response to *A. mellifera* queen pheromone, and *D. melanogaster* and *Bombus terrestris* pheromone-response to *B. terrestris* queen pheromone. Number indicates the number of genes associated with each module.

**Table S1.** *Drosophila melanogaster* ovary size by pheromone treatment. Flies were exposed to treatment pheromone from egg until collection. Virgin female flies were collected and dissected five days after emergence. SD denotes standard deviation.

| Treatment | Replicates | Mean size (mm<sup>2</sup>) | SD |
| --- | --- | --- | --- |
| Bumblebee | 10 | 0.32 | 0.064 |
| Termite | 10 | 0.35 | 0.145 |
| QMP | 10 | 0.51 | 0.085 |
| QMP and Brood | 10 | 0.59 | 0.156 |
| Brood | 10 | 0.65 | 0.123 |
| Control (2nd) | 10 | 0.80 | 0.146 |
| Control (1st) | 10 | 0.88 | 0.136 |

**Table S2.** Multiple Comparisons of Means: Tukey Contrasts for ovary sizes of each pheromone treatment. Highlighted comparisons are statistically significantly different. All treatments are significantly different to the controls. The controls are not significantly different.

| Group 1 | Group 2 | Estimate | p.Val |
| --- | --- | --- | --- |
| Termite | Bumblebee | 0.027 | 1.000000 |
| QMP | Bumblebee | 0.189 | 0.014997 |
| QMP and Brood | Bumblebee | 0.270 | 0.000026 |
| Brood | Bumblebee | 0.328 | 0.000000 |
| Control (2nd) | Bumblebee | 0.483 | 0.000000 |
| QMP | Termite | 0.162 | 0.076808 |
| QMP and Brood | Termite | 0.244 | 0.000262 |
| Brood | Termite | 0.301 | 0.000002 |
| Control (2nd) | Termite | 0.456 | 0.000000 |
| QMP and Brood | QMP | 0.082 | 1.000000 |
| Brood | QMP | 0.139 | 0.262967 |
| Control (1st) | QMP | 0.371 | 0.000000 |
| Brood | QMP and Brood | 0.058 | 1.000000 |
| Control (1st) | QMP and Brood | 0.289 | 0.000005 |
| Control (1st) | Brood | 0.231 | 0.000823 |
| Control (1st) | Control (2nd) | 0.076 | 1.000000 |

## Notes

### Competing Interest Statement

The authors have declared no competing interest.

### Summary of Updates

Considerable reworking of text, analysis pipeline and figures.

